# Blinatumomab-driven T-cell activation in αβ and γδ T-cell subsets: Insights from in vitro assays§

**DOI:** 10.1101/2025.11.17.688178

**Authors:** Miriam Kelm, Nourhan Nasr, Sonja Bendig, Dieter Kabelitz, Marta Lustig, Heiko Trautmann, Anna Laqua, Christian Peters, Daniela Wesch, Heiner Oberg, Ottmar Janssen, Thomas Valerius, Claudia D Baldus, Alexander Georg Scheffold, Monika Brüggemann, Guranda Chitadze

## Abstract

Blinatumomab (BLN) is a bispecific T-cell engager that has revolutionized the treatment of B-cell precursor acute lymphoblastic leukemia (BCP-ALL), significantly improving outcomes in both adults and children. By simultaneously binding to CD19 on B cells and CD3 on T cells, BLN triggers target cell-dependent T-cell activation, resulting in the cytolysis of CD19^+^ BCP-ALL cells. Despite the remarkable clinical advancements achieved with BLN, the immunological mechanisms underlying treatment response or failure remain poorly characterized. γδ T cells are attractive candidates for adoptive T–cell therapy due to potent cytotoxicity, capacity to present antigens, broad lysis of different tumor entities, and low alloreactivity. Because γδ T cells can also be redirected by BLN, we systematically studied BLN–driven effector functions of conventional αβ and unconventional γδ T cells.

We evaluated cytotoxicity and cytokine/effector release in freshly isolated and in vitro-expanded αβ and γδ T cells from healthy adults against CD19⁺ BCP-ALL lines (NALM-6, HAL-01), and profiled dynamic phenotypic alterations by multiparametric flow cytometry. CD19⁺ targets were consistently reduced in the presence of BLN. Freshly isolated αβ, especially CD8⁺, displayed superior BLN-mediated cytotoxicity as compared to γδ T cells, with donor-dependent variability in γδ killing. Notably, zoledronate-expanded Vγ9Vδ2 γδ T-cell lines achieved cytotoxicity comparable to PHA-expanded αβ cells. However, γδ T-cell-killing benefited from higher BLN concentration when challenged with high tumor load. BLN induced CD3 down-modulation in αβ T cells but not in γδ T cells, alongside higher soluble Fas ligand in αβ cultures, consistent with stronger early activation, preceding activation-induced cell death. γδ T cells showed no such changes, suggesting reduced susceptibility to activation-induced cell death. Single-cell RNA and flow analyses corroborated these findings, showing robust activation/exhaustion programs in αβ T cells and a stable effector-memory state with low checkpoint expression in γδ T cells.

Together, these data reveal subset–specific BLN responses and support expanded Vγ9Vδ2 γδ T cells as a rational adoptive partner to BLN — particularly in settings of favorable antigen density/low tumor burden — providing complementary cytotoxicity with potentially reduced inflammatory liability. These findings provide a framework for combining γδ T-cell-based therapies in BLN-treated patients for improving BLN efficacy in BCP-ALL patients.

## Introduction

The outcome of B-cell precursor acute lymphoblastic leukemia (BCP-ALL) has improved significantly over the last decade due to personalized therapy protocols and novel therapeutic strategies, including mono- or bispecific antibodies, antibody-drug conjugates, and genetically modified CAR T cells^1,2^. Nevertheless, relapse remains common, and more than 20% of patients still succumb to the disease. Outcomes are notably poorer in adults than in children, with five-year overall survival rates of around 90% in pediatric cohorts but only 40–60% in adults^3–6^. This highlights the critical need to understand the mechanisms underlying therapy failure and to further optimize clinical outcomes.

Blinatumomab (BLN), a bispecific T-cell engager linking CD3 on T cells with CD19 on B cells, facilitates the formation of immunological synapses, leading to T-cell activation, expansion, and targeted lysis of CD19⁺ cells^2,7^. Since its initial FDA approval in 2017 for relapsed or refractory B-cell precursor ALL (R/R BCP-ALL) in both adults and children, BLN has demonstrated remarkable clinical efficacy^8–11^. Nonetheless, many patients relapse or fail to achieve long-term remission after BLN therapy in R/R or frontline in minimal residual disease (MRD) settings^9,12,13^. Approximately 25% of these failures are due to loss of target antigen; the rest of the relapse cases however are due to impaired T-cell functionality^14^. In heavily pretreated patients, such dysfunction may arise from prior chemotherapy regimens ^15^, from high leukemic burden^8,13^ or from persistent T-cell stimulation due to BLN’s administration as a 28-day continuous infusion, required due to its short half-life^16,17^.

Given these limitations, exploring novel strategies to enhance BLN-mediated cytotoxicity through complementary therapeutic approaches is essential. One promising approach involves the transfer of γδ T cells from unrelated donors, a minor yet highly potent subset of T cells characterized by CD3-associated TCR γδ expression and intrinsic anti-tumor cytotoxicity irrespective of HLA-matching^18–20^. Adoptive γδ T-cell therapy is attractive because γδ T cells recognize tumor cells through stress-induced ligands without relying on HLA presentation. γδ T cells have a reduced risk of graft-versus-host disease (GvHD) and are suitable for allogeneic application across diverse patient populations^21–23^.

Peripheral γδ T cells constitute up to 5-10% of circulating T cells, predominantly represented by the Vγ9Vδ2 γδ T-cell subset^18,24,25^. These cells respond robustly to pyrophosphates (“phosphoantigens”) frequently accumulating in tumor cells, and their selective activation and expansion can be reliably induced using zoledronic acid (Zole) or synthetic phosphoantigens, thus facilitating clinical-grade ex vivo expansion^26,27^. γδ T-cell activation depends on interactions with BTN3A1 and BTN2A1^28,29^. Additionally, γδ T cells can function as antigen-presenting cells, cross-presenting antigens to αβ T cells and further enhancing immune responses^30,31^. Their broad tumor-recognition capacity, combined with independence of HLA-restriction, positions γδ T cells as highly favorable candidates for adoptive cell therapies across diverse malignancies, as currently explored in numerous clinical studies utilizing expanded Vδ1 and Vδ2 T-cell subsets^19^. Clinical safety of allogeneic Vγ9Vδ2 γδ T-cell immunotherapy has already been documented, with prolonged survival reported in patients with late-stage lung or liver cancer^32^.

Our own results suggest prognostic value of γδ T cells in ALL, with increased levels correlating to improved MRD responses post-chemotherapy^33^. Limited preclinical evidence, supports the therapeutic potential of combining adoptively transferred γδ T cells with CD3-engaging antibodies, including BLN^34^. However, their ability to augment BLN-mediated cytotoxicity in adoptive immunotherapy settings has not been fully evaluated. Moreover, γδ T cells may follow distinct exhaustion pathways compared to αβ T cells^35,36^, but this has not been investigated in the context of BLN.

In this study, we systematically investigated BLN-mediated changes in the effector functions of ex vivo expanded γδ T cells compared to αβ T cells. We assessed their phenotypic and functional fitness under conditions mimicking variable tumor burdens typically encountered at diagnosis or during therapy. Our approach combined functional assays, multiparametric phenotyping, and single-cell transcriptomics to dissect how γδ T cells kill CD19⁺ targets in the presence of BLN. A deeper understanding of these mechanisms may inform novel therapeutic strategies to improve outcomes of BLN-treated patients with BCP-ALL.

## Materials and Methods

### Cell cultures

Peripheral blood mononuclear cells (PBMCs) were isolated from leukocyte concentrates of healthy adult blood donors obtained from the Institute of Transfusion Medicine (Institute for Transfusion Medicine, University Hospital Schleswig-Holstein Campus Kiel) using Ficoll density gradient centrifugation (Merck, Darmstadt, Germany). Ethical approval was granted by the institutional ethics review board of the University Medical Faculty Kiel (approval number D405/10, D479/18, D546/16). The research was conducted in accordance with the Declaration of Helsinki. CD4⁺, CD8⁺, and γδ T cells or when required entire CD3^+^ T cells were purified from PBMCs by magnetic-activated cell sorting (MACS) using respective negative selection kits according to the manufacturer’s instructions (Miltenyi Biotec, Bergisch Gladbach, Germany). To establish αβ T-cell lines, PBMCs were stimulated with Phytohemagglutinin A (PHA; 0.5 µg/mL; Thermo Fisher Scientific, Waltham, MA, USA) and recombinant interleukin-2 (rIL-2; 100 U/mL; Novartis, Basel, Switzerland) in culture medium which consists RPMI 1640 medium (Thermo Fisher Scientific) supplemented with penicillin (100 U/mL), streptomycin (100 µg/mL), and 10% heat-inactivated fetal bovine serum (FBS; Thermo Fisher Scientific), hereafter referred to as culture medium, following previously established protocols^37,38^. Vδ2 T cells were expanded using Zoledronate (Zole, 2.5 µM; Novartis) and rIL-2 (50 IU/mL) to establish short-term activated Vδ2 T-cell lines as previously reported^39^. IL-2 was added every 2–3 days and cultures were split after day 6 or 7. After 14 days, non-viable cells were removed via Ficoll density centrifugation when necessary. The purity of expanded cell lines was assessed at day 14 by flow cytometry.

The human B-cell precursor ALL cell lines NALM-6 and HAL-01 obtained from DSMZ and maintained in required culture medium according to manufacturer’s instructions and utilized as target cells in co-culture assays.

### Co-culture experiments

Freshly isolated CD4⁺, CD8⁺, and γδ T cells were co-cultured with HAL-01 cells in culture medium, with or without BLN (20 ng/mL) for up to 7 days^40^. T-cell numbers were assessed via manual counting at day 3 and day 7 and analyzed for their composition and phenotypes by flow cytometry at day 3. Supernatants were collected at specified day 3 and day 7, stored at −20 °C, and analyzed via ELISA as described below. In vitro expanded PHA-activated αβ and Zole-activated γδ T cells were co-cultured with NALM-6 or HAL-01 target cells at various effector-to-target (E:T) ratios in the presence or absence of BLN (20 ng/mL or 0.5 ng/mL) for 24 hours or 3 days. PBMCs from 6 healthy donors were co-cultured for 7 days with the CD19⁺ BCP-ALL cell line HAL-01 at different effector-to-target (E:T) ratios: 1:1 (high CD19 load), 5:1 (medium load), or without addition of exogenous target cells (low CD19 load, autologous B cells only), in the presence of in the absence of BLN (20 ng/mL). Cells were harvested and analyzed on days 0, 3, and 7 by multiparametric flow cytometry. Effector cells at fixed numbers were cultured with varying amounts of target cells through all co-culture experiments. Recombinant human IL-2 (50 IU/mL) was added to all co-cultures at baseline and replenished every 48 h (no other exogenous cytokines were used). Media volume lost to sampling was replaced with fresh complete medium containing the same IL-2 concentration.

### Multiparametric flow cytometry

Purity of PHA- and Zole-expanded αβ and γδ T-cell cultures and MACS-isolated T-cell populations were assessed by flow cytometry using anti-CD3, CD4, CD8, γδ TCR, and αβ TCR monoclonal antibodies. Only populations with >80-90% purity were used in co-culture assays (Supplemental Table 1, Supplemental Figure 1).

B-cell depletion in BLN-activated PBMCs or in co-cultures was analyzed on BD FACS Canto using antibodies directed against CD19 and CD3 or at Cytek Northern Lights cytometer using a 22-color antibody panel also incorporating anti-CD3 and anti-CD19 with the following antibodies directed against CD4, CD8, CD25, CD27, CD28, CD38, CD45, CD45RA, CD56, CD95, Annexin V, CCR7, DNAM-1, HLA-DR, PD-1, γδ TCR, TIGIT, TIM-3, Vδ2 γδ TCR, and viability dye 7-AAD.

Selected samples undergoing single-cell RNA sequencing (see below) were analyzed using antibodies against CD3, CD4, CD8, CD16, CD19, CD22, CD24, CD45, CD45RA, CD56, CCR7, DNAM-1, γδ TCR and Vδ2 γδ TCR. Samples were analyzed on a BD FACSLyric. Detailed information of used antibody panels and used clones or formats is provided in Supplemental Table 2.

### Enzyme-linked immunosorbent assay (ELISA)

Cell culture supernatants were analyzed for Interferon-γ (IFNγ; DIF50C), Granzyme B (GrzB; DGZB00), Perforin (QK8011), Granulysin (Gnly; DY3138), Perforin (Prf; QK8011) and soluble Fas-Ligand (sFAS-L; DFL00B) using sandwich ELISA kits from R&D Systems, Minneapolis, USA according to the manufacturer’s instructions. Absorbance was measured at 450 nm using a microplate reader (Tecan Group Ltd, Männedorf, Switzerland) in the Institute of Immunology Kiel.

### Determination of CD19 Antigen-density

To quantify CD19 antigen density on the cell surface of BCP ALL cell lines (NALM6 and HAL-01), the specific antibody binding capacity (SABC) was determined using the purified anti-human CD19 antibody (clone HIB19) and the QIFIKIT® (DAKO, Glostrup, DK) by flow cytometry according to the manufacturer’s instructions in the Division of Stem Cell Transplantation and Cellular Immunotherapies.

### BD Rhapsody single-cell RNA-seq

BD Rhapsody Single-Cell Analysis System (BD, Biosciences) were utilized using a targeted approach with human T-cell Expression Panel commercially available from BD Biosciences and covering 259 genes, commonly expressed in human T cells (Cat ID 633751, Supplemental Table 3). For this, PBMCs from a healthy donor were cultured for 72 hours with rIL-2 (50 IU/mL) + BLN (20 ng/mL) or with rIL-2 alone as a control. Afterwards, CD3⁺ cells were negatively isolated (human MACS Pan T cell isolation Kit, Miltenyi Biotec), labeled using BD Single-Cell Multiplexing Kit and AbSeq (antibody-labeled with oligos) reagents targeting CD4, CD8, CD25, CD45RA, CD127, CCR7, and TCRγδ (Supplemental Table 2). Cells from BLN-treated and control samples were labeled with Sample Tags following the manufacturer’s instructions (Single Cell Labelling with the BD™ Single-Cell Multiplexing Kit and BD™ AbSeq Ab-Oligos (ID: 214419 Rev. 2.0)). 10,000 single cells were load on a cartridge with preloaded beads to capture RNA of single cell, followed by cDNA synthesis using the BD Rhapsody Express Instrument and Scanner following BD protocols (Single Cell Capture and cDNA Synthesis with the BD Rhapsody™ Single-Cell Analysis System (210966 Rev. 1.0)). cDNA libraries of target transcripts, Samples Tags and Abseqs were prepared using the Manufactures instructions, following mRNA Targeted, Sample Tag, and BD™ AbSeq Library Preparation with the BD Rhapsody™ Targeted mRNA and AbSeq Amplification Kit (214508 Rev. 3). The final libraries were quantified using a Qubit Fluorometer with the Qubit dsDNA HS Kit (ThermoFisher) and the size-distribution was measured using the Agilent high sensitivity D5000 assay on a TapeStation 4200 system (Agilent Technologies). Sequencing was performed in paired-end mode (2×75 cycles) on NextSeq 500 System (Illumina) with NextSeq 500/550 Mid Output Kit, generating approximately 1.3 billion reads. Data processing utilized the Seven bridges pipeline BD Rhapsody™ Targeted Analysis Pipeline - Revision: 0. Generated demultiplexed matrices of ScRNA-seq UMI count were imported to R 4.3.3 and gene expression data analysis was performed using the R/Seurat package 5.3.0^41^. To remove doublets and cell fragments, we further fitted a linear model of detected genes versus transcript count per sample and excluded deviating outliers (0.5 – 5 % of cells, depending on sample quality). Additionally, we removed genes that were expressed in less than ten cells. Before downstream analysis, LogNormalization (Seurat function) was applied separately to the RNA and AbSeq assays. The data were then scaled regressing for total UMI counts and principal component analysis (PCA) was performed. RNA and AbSeq data were integrated using weighted nearest neighbors (WNN) based on the first 8 RNA and first 6 AbSeq principal components. UMAP and clustering were performed on the resulting WNN graph. For two-dimensional data visualization we performed UMAP based on the first 20 dimensions of the resulting WNN graph. The cells were clustered using the Louvain algorithm based on the first 20 dimensions with a resolution of 0.8.

### Statistical analysis

For statistical comparisons of paired data, we used two-tailed paired T-tests. For unpaired data and comparing independent variables we used Mann-Whitney U-tests. Statistical significance was set at p<0.05, denoted as follows: p<0.05 (*), p<0.01 (**), p<0.001 (***). Analyses were performed using GraphPad Prism Software 8.4.3 and R-Studio 4.3.3 software.

## Results

### BLN potentiates cytotoxicity of fresh αβ and γδ T cells, but αβ T cells dominate early cytotoxicity

We first evaluated BLN-induced killing of CD19⁺ target cells by freshly isolated γδ, CD4⁺ and CD8⁺ αβ T-cell populations. Isolated T-cell populations (γδ, CD4⁺ and CD8⁺ αβ) were co-cultured with the CD19⁺ malignant BCP ALL cell line HAL-01 at an effector-to-target ratio of 5:1 in the presence of BLN at concentration of 20 ng/mL, previously demonstrated to effectively induce immunological synapse formation^40^. At day three, target CD19⁺ cells were consistently reduced in all co-cultures in the presence of BLN, as measured by assessing residual CD19^+^ cells using flow cytometry (Figure 1A, top panel shown for CD8^+^ αβ T cells). However, γδ T cells displayed lower cytolytic activity compared to CD8^+^ and CD4^+^ αβ T cells, demonstrated by persistence of CD19^+^ target cells in γδ and not in αβ T-cell cultures in the presence of BLN (Figure 1A, bottom panel).

**Figure 1:**
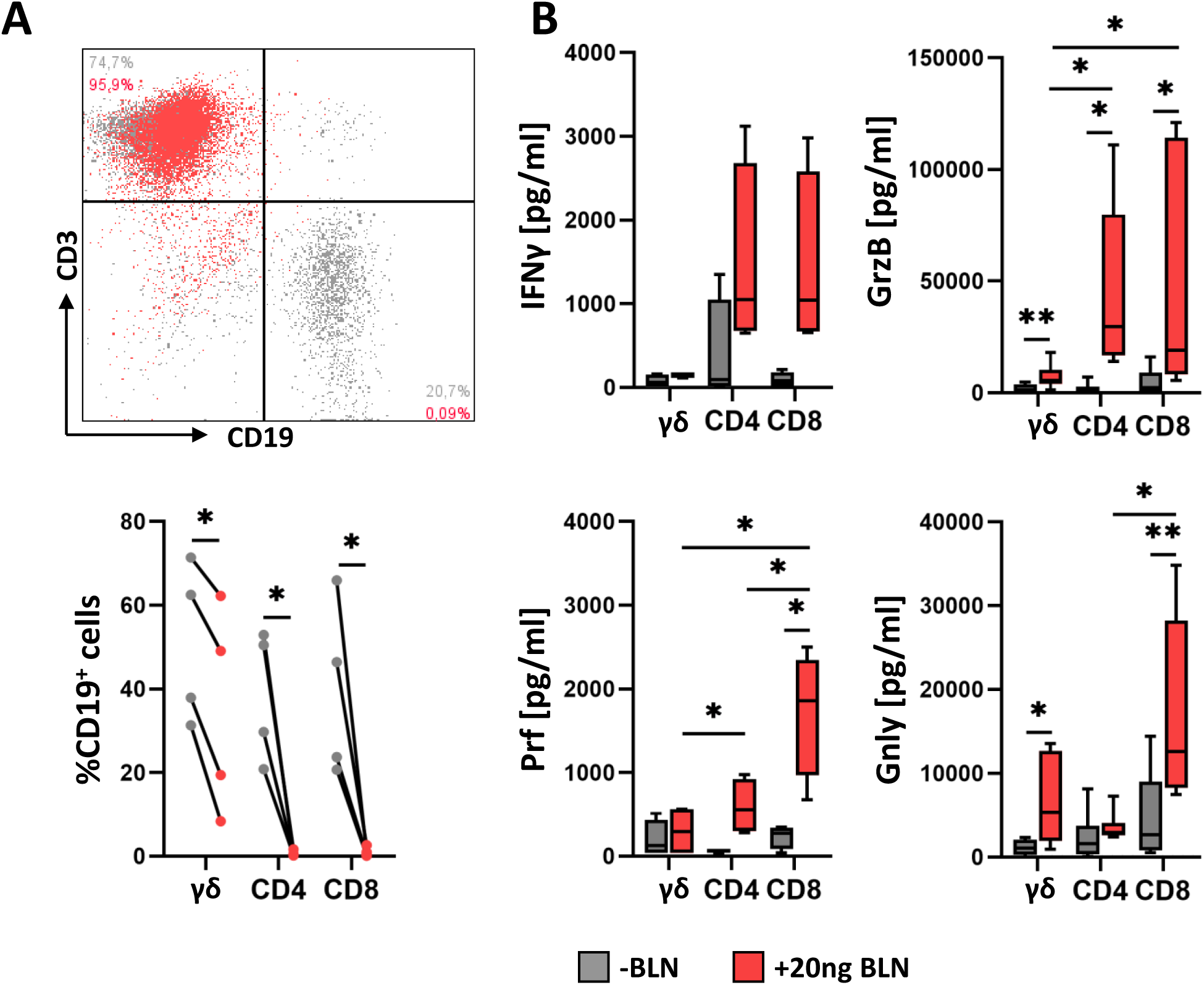
BLN-mediated T-cell responses against CD19+ malignant B-cell line. A: BCP-ALL cells in the presence (red) or absence (grey) of BLN for three days were co-cultured with freshly isolated γδ, CD4⁺ and CD8⁺ αβ T cells/ T- and B-cell frequencies in the coculture were assessed by staining for CD3 and CD19, as exemplary shown for CD8⁺ αβ T cells. B: The release of IFNγ, GrzB and Prf into supernatants of cocultures in the presence (red) or absence (grey) of BLN was assessed via ELISA after three days, and the release of Gnly was analyzed after seven days. Data includes 4 to 7 healthy donors and * indicates p value between 0.01-0.05, ** indicates p value between 0.01-0.001 (paired and unpaired T-test).

ELISA assays revealed significantly increased secretion of IFNγ and cytotoxic effector molecule Granzyme B (GrzB), Perforin (Prf) and late mediator of cytotoxicity Granulysin (Gnly)^42^. At day 3 in BLN-treated cultures, with αβ T cells releasing consistently higher levels than γδ T cells. Prf levels were highest in CD8⁺ T-cell cultures at day 3, whereas Gnly levels, assessed after seven days, were significantly elevated in both CD8⁺ and γδ T-cell cultures treated with BLN (Figure 1B).

Overall, BLN enhanced cytotoxic potential of freshly isolated γδ and αβ T-cell populations. However αβ T-cell populations demonstrated superior BLN-mediated cytolytic activity compared to γδ T cells, with CD8⁺ T cells having the potentially highest cytotoxic potential, as indirectly indicated by perforin secretion^43^.

### Activated γδ T-cells match αβ cytotoxicity under favorable antigen conditions

Given the known cytotoxic potential of Zole-activated Vδ2 T cells and their potential clinical application^19,44^, we used same read out and assessed residual CD19^+^ in flow-based assay following coculture with Zole-activated Vδ2 γδ T-cell lines instead of freshly isolated γδ T cells (Figure 2A). We studied release of cytokine mediators/effector molecules at 24 hours anticipating stronger effector functions (Figure 2B) in the presence of BLN by Zole-expanded γδ T cells (Figure 2A, green) and compared it with effector functions mediated by Phytohemagglutinin (PHA)-expanded αβ T cells (Figure 2A, grey) against two BCP-ALL cell lines, HAL-01 and NALM-6, with high and low expression CD19-patterns respectively (Supplemental Figure 2). At an effector-to-target ratio of 5:1 and BLN concentration of 20 ng/mL, both γδ and αβ activated cultures resulted in significant CD19^+^ reduction. However, PHA-expanded αβ T cells induced a slightly more pronounced B-cell killing with complete clearance of CD19^+^ cells. Target cells still persisted in Zole-expanded γδ T cells after 24 hours at low levels, especially in a case of NALM6 (residual CD19^+^ cells at median value of 4.4% of co-culture), expressing lower levels of CD19 compared to HAL-01 (Supplemental Figure 2). Interestingly, we observed CD3 downregulation, that selectively occurred in PHA-expanded αβ T cells in the presence of BLN, whereas Zole-expanded γδ T cells maintained stable CD3 expression (Figure 2A, bottom panel). BLN-independent T-cell responses were also observed, as indicated by partial B-cell elimination in the absence of BLN occurring in a donor-dependent manner, with PHA-expanded αβ T cells (dark grey) outperforming Zole-expanded γδ T cells (dark green).

**Figure 2:**
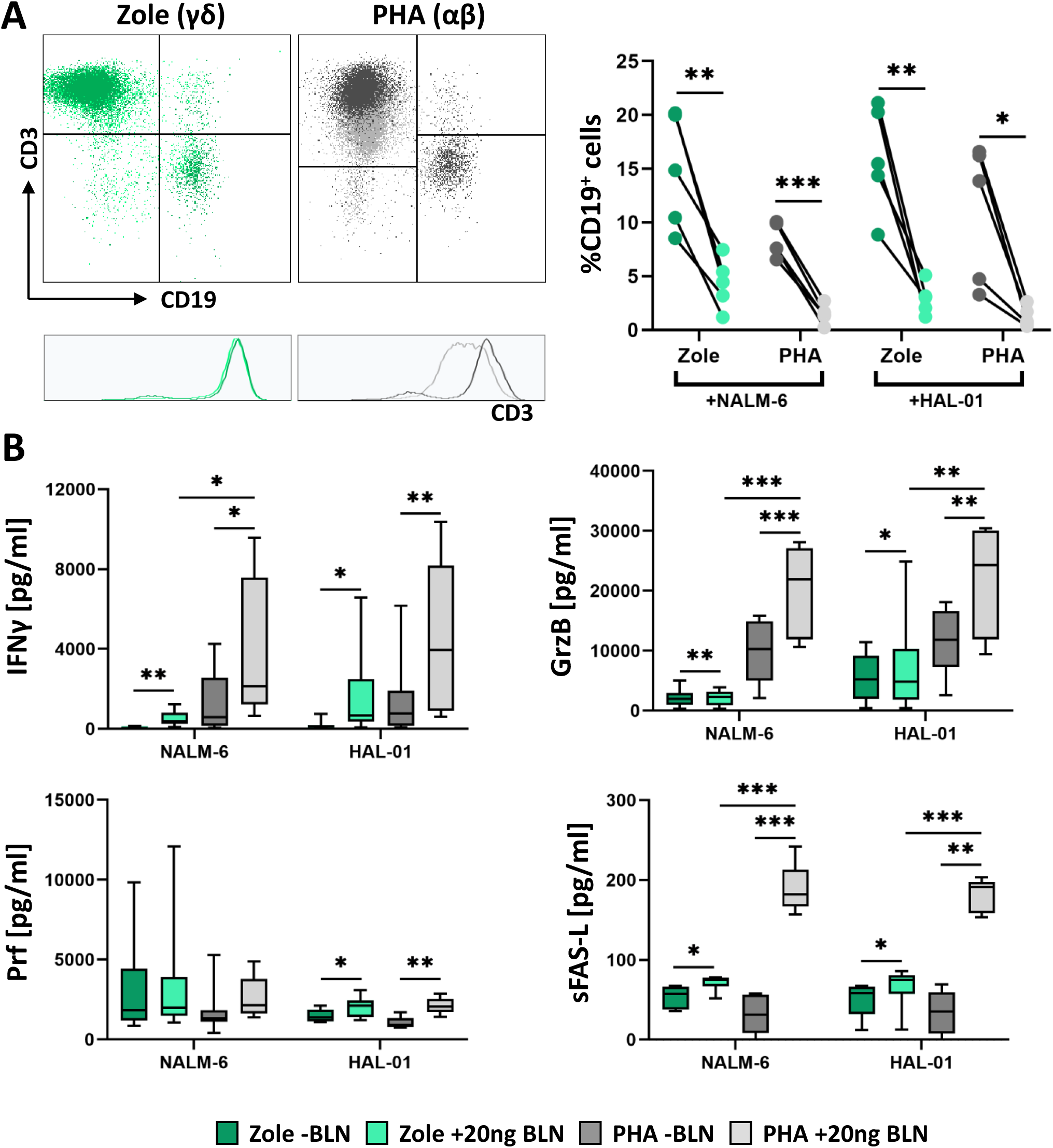
CD19^+^ target cell elimination by activated αβ and γδ T-cell cultures. In vitro PHA-expanded aβ (grey) and Zole-expanded γδ T cells (green) were cocultured with different malignant B-cell lines (NALM-6, HAL-01) in the presence or absence of BLN for 24 h. A: Exemplary staining at 24h, defining residual CD19^+^ cells and CD3^+^ in the co-culture in presence of BLN and PHA or Zole activated T cells. B: The release of IFNγ, GrzB, Prf and sFAS-L was analyzed in respective coculture supernatants of BLN-treated or untreated cells via ELISA. Data includes 4 to 7 healthy donors and * indicates p value between 0.01-0.05, ** indicates p value between 0.01-0.001, , *** indicates p value between < 0.0001 with paired T-test.

Both expanded γδ and αβ T-cell lines exhibited increased secretion of IFNγ, GrzB, soluble FAS ligand (sFAS-L), and Prf following BLN stimulation in the presence of tumor cells. IFNγ and GrzB were lower in Zole-expanded cultures compared to PHA-expanded cultures (Figure 2B). Interestingly, high levels of sFAS-L, known to occur during activation induced cell death (AICD) expected in case of excess of T-cell activation^45,46^, was higher in the presence of PHA-expanded αβ T cells compared to Zole-expanded γδ T cells. Thus, when comparing Zole and PHA-activated T-cell cultures, αβ T-cells exhibit superior cytotoxicity; however, release higher levels of soluble FAS-L and potentially undergo CD3 downregulation in contrast to γδ T cells.

### Target antigen density/tumor burden jointly impact γδ efficacy; higher BLN dose preferentially benefits γδ under stress

In clinical setting, response to BLN is largely dependent on ALL blast counts^8^. Blinatumomab has higher affinity to CD19 and lower affinity to CD3^47^. BLN binds to CD19 and acts as activation matrix for T cells, that can detach and kill up to 5-10 target cells within 9 hours (Serial killing)^48,49^. In the absence of CD19^+^, no T-cell activation takes places^47^. Thus, CD19 target expression is relevant for BLN mediated T-cell activation and target antigen density/target cell load, as well as the concentration of BLN might impact T-cell signaling and activation^8^. Thus, we next explored how varying the effector-to-target ratio and BLN concentration influenced γδ T-cell cytotoxicity. Using the HAL-01 cell line, which exhibits high CD19 antigen density (Supplemental Figure 2), we cultured Zole-expanded γδ T-cell lines at 5:1 (experimental settings from Figure 2) and 1:5 effector-to-target ratios, corresponding to low and high ALL load, respectively (Figure 3). BLN was tested at 20 ng/mL and at concentration of 0.5 ng/mL, close to steady-state serum level reported in patients^50^. Residual target (CD19^+^) cells and T-cell composition/phenotypes linked with T-cell functionality were evaluated after 24 hours and following three days by 22-color spectral flow cytometry (Supplemental Table 2).

**Figure 3:**
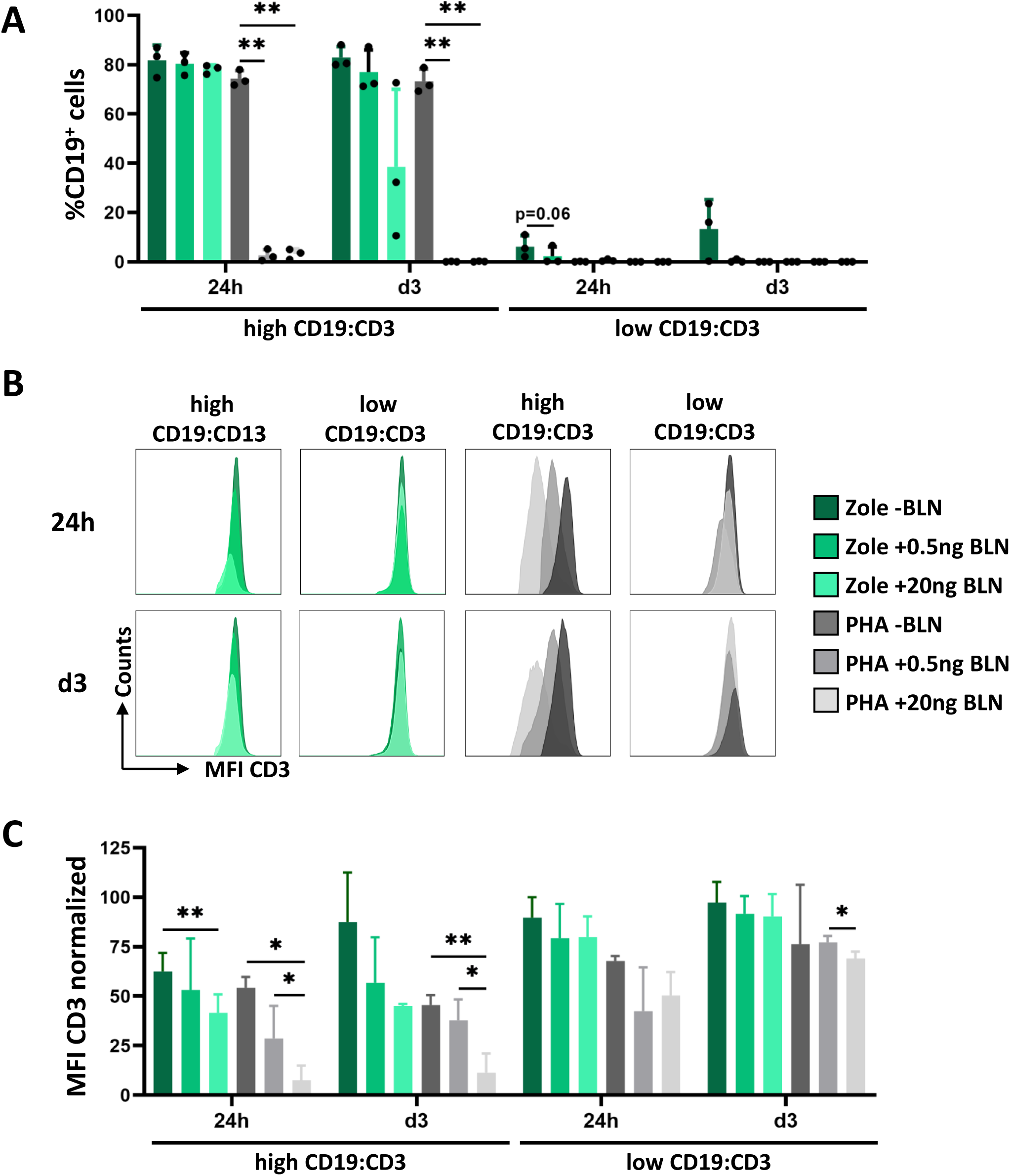
Impact of tumor burden and BLN concentration on T-cell-mediated cytotoxicity in Zole (γδ) - and PHA (αβ)-expanded T cells. A: Median B-cell decline of HAL-01 cells from Zole-(green) and PHA-expanded T cells (grey). Individual values are represented by dots. B: Representative example of CD3-expression in γδ and αβ T-cell cultures, following BLN/mediated activation. C: Normalized CD3 MFI of different culture conditions. High T = High Tumor load (E:T 1:5), Low T = Low Tumor load (ET 5:1). CD3 was normalized to unstained control within the same experiment. N=3. Data includes 3 healthy donors and * indicates p value between 0.01-0.05, ** indicates p value between 0.01-0.001 with paired T-test.

Both γδ (Zole-expanded, green) and αβ (PHA-expanded, grey) T-cell cultures efficiently eliminated target cells under low tumor burden conditions (Figure 3A, right panel, low CD19:CD3, similar to data shown in Figure 2). At high CD19:CD3 conditions (Figure 3A, left panel), αβ T-cell cytotoxicity was already maximal at 24 hours and did not significantly vary with BLN concentration. However, BLN did not enhance γδ T-cell mediated killing at low BLN concentrations. Higher BLN concentrations were more beneficial to improve γδ T-cell cytotoxicity, particularly at day 3 in case of two donors (Figure 3A, left panel, grey bars). Consistent with previous findings, marked CD3 downregulation was observed predominantly in αβ T cells upon BLN treatment, especially at high tumor burden conditions (Figure 3B-C; Supplemental Figure 3). Because CD3 down–modulation occurred only at high target burden, and spared γδ T cells while primarily affecting αβ T cells under identical BLN and staining conditions, we interpret this as CD3 internalization after strong T–cell stimulation rather than epitope masking^51^.

### Zole–expanded γδ T cells retain a stable effector memory (EM) phenotype with low checkpoint expression under BLN

The differential CD3 modulation observed in γδ and αβ T cells suggested distinct BLN-mediated signaling through CD3, which we hypothesized would result in phenotypic and functional differences, affecting their fitness and potentially susceptibility to AICD, as supported by the high levels of sFAS-L detected in αβ T-cell cultures. Of note, sFas-L reflects ADAM10/17–mediated shedding of the pro–apoptotic membrane FasL and elevated sFasL in CD8⁺ cultures are here interpreted as a biomarker of strong re-stimulation and shedding, not as a direct driver of AICD^45,46,52^.

To investigate this, we compared the composition and phenotype of Zole- and PHA-expanded T cells at baseline (prior to co-culture) and analyzed their numbers and phenotypic changes at day 1 and day 3 using spectral flow cytometry (Figure 4). At baseline prior co-culture (d0), after 14 days of expansion, T-cell composition differed markedly: Zole-expanded cultures consisted expectedly of Vδ2 γδ T cells (median 90%, range 80-96%), while PHA cultures comprised both CD4⁺ and CD8⁺ αβ T cells with median proportions of 27% (range 13%-31%) and 65% (range 63%-83%), respectively (Supplemental Table 1).

**Figure 4:**
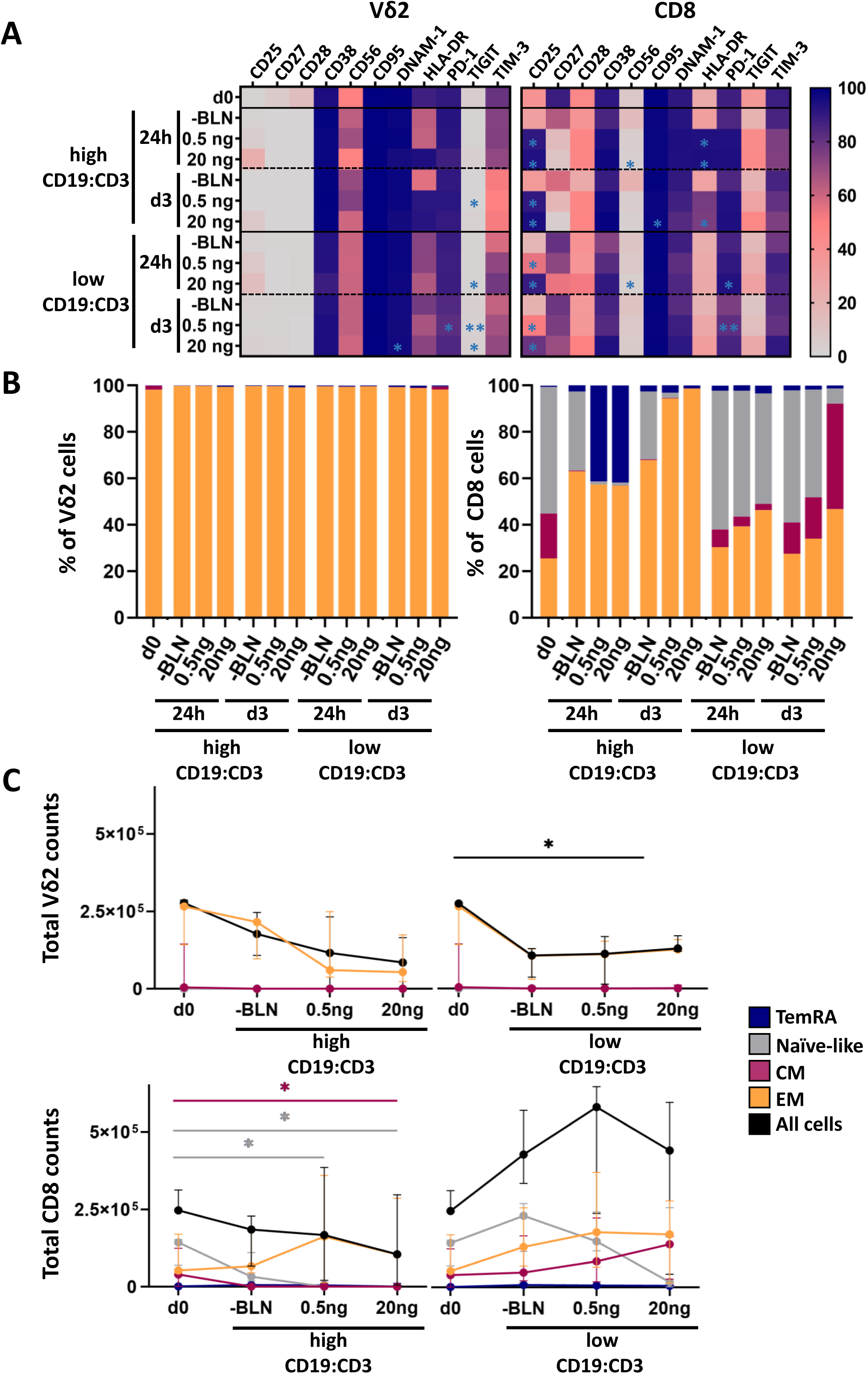
Phenotypic differences between activated γδ and αβ T cells. A: Median Marker Expression on Zole-expanded Vδ2 and PHA-expanded CD8 T cells of three healthy donors. B: Median values of EM, CM, Naïve and TEMRA cells of CD8 and Vδ2 cells. B: Absolute cell counts of EM, CM, Naïve and TEMRA cells of CD8 and Vδ2 cells at day 0 and day 3. Error bars representing Median value with 95% confidence interval. Stars representing significant changes between baseline and BLN therapy at day 1 and day 3.

Zole-expanded Vδ2 γδ T cells displayed a highly homogeneous effector memory (EM) phenotype (CD27⁻CD45RA⁻), consistent with previous observations^39,44^. Their absolute cell counts did not significantly change during three days of co-culture across different BLN concentrations at high ALL load (Figure 4B, black line). We observed slight decrease in absolute counts in low CD19:CD3 conditions (Figure 4A, top panel). Day 14 Zole-expanded γδ T cells expressed high levels of activation markers (CD38, HLA-DR, DNAM-1, CD56) while showing lower or absent levels of exhaustion markers such as TIGIT (baseline, d0). During co-culture there was not significant change of their phenotype compared to baseline (Figure 4A). TIGIT negativity on day 14 cultures goes in line with a reported transient peak around day 4 and declining by day 7 in Zole-expanded T-cell cultures.^35^

In contrast, PHA-expanded CD8⁺ αβ T cells were more heterogeneous with naïve and memory subsets at baseline. Upon BLN stimulation, especially in the context of high tumor load, naïve cells declined and EM and TemRA (CD27^-^CD45RA^+^) subsets increased after 24 hours, with EM cells becoming dominant by day 3 (Figure 4A). This was reflected by an increase in total EM counts in the presence of high tumor load at day 3, as single dominant population. In cultures with lower CD19⁺ burden, in line with less activation provided by low target cell density, naïve CD8⁺ T cells transitioned toward both EM and central memory (CM) phenotypes. Interestingly absolute counts in the presence of low target cells and lower BLN amount was higher compared to high target density and high BLN concentration. Phenotypic shifts observed in PHA-expanded CD4⁺ αβ T cells following BLN are shown as supplemental figure 4.

Phenotypically, PHA-expanded CD8⁺ αβ T cells showed strong activation in response to BLN, characterized by increased expression of CD25 and HLA-DR in line to observed differentiation from naïve to memory phenotypes. TIGIT expression was markedly high in αβ T cells at baseline. We also attempted to study differences in AICD at 24h or at day 3, however we did not observe differences in annexin–V expression on T cells following BLN stimulation (readout for AICD), possibly because AICD often peaks within 4–12 h of stimulation.

### BLN expands CD4⁺ and CD8⁺ but not γδ T cells from mixed PBMC cultures

Next, we asked whether BLN treatment in a mixed PBMC context could induce γδ T-cell proliferation alongside αβ T cells, despite unfavorable abundances of γδ T cells. This particular point is important in the clinical setting. To this end we stimulated PBMCs together with CD19^+^ HAL-01, at an E:T ratio of 1:1 (corresponding more to high CD19^+^ load when taking CD3 T-cell proportions of PBMCs into account), at an E:T ratio of 5:1 (medium CD19^+^ load) and without addition of CD19-expressing cell lines (low CD19^+^ load, autologous B-cells) for 7 days with (red) and without (grey) 20 ng/ml BLN (Figure 5A). We assessed absolute cell counts of CD4^+^, CD8^+^ and γδ T cells at day 0, day 3 and day 7 of the co-culture and assessed immune profiles of T cells. We noticed a significant expansion of CD4^+^ and CD8^+^ T cells in all three culture conditions with BLN treatment with low, medium and high CD19^+^ load. In contrast to CD4^+^ and CD8^+^ T cells, we did not see a noteworthy expansion of γδ T cells under BLN treatment from PBMCs up to day 7. As expected there was no significant increase in T-cell counts in the absence of BLN (Figure 5, grey lines). Interestingly we saw higher expansion rates of both CD4^+^ and CD8^+^ T cells in the presence of high target cell density compared to other two conditions with medium and low target cell density at day 3. At day 7, there were no differences among these conditions in CD4^+^ T cells. In CD8^+^ T cells however, the cultures with the highest CD19^+^ cells at baseline had the lowest CD8^+^ T-cell counts at day 7 compared to the other two cultures with medium and low frequencies of CD19^+^ cells (red filled mark). This was confirmed by strongest activation profile (CD25^+^, HLA-DR^+^) in the presence of high target cells (not shown), compared to other cultures, where EM CD8^+^ T cells predominated the BLN-containing cocultures on day 7, whereas the “naïve” population almost disappeared. In summary BLN application did not lead to γδ T-cell expansion in the presence of other T-cell populations, CD8⁺ T cells were the most expanded subset, as previously reported^53^ and target load was inversely corelated with absolute CD8+ T-cell counts, suggestive of AICD following strong activation.

**Figure 5:**
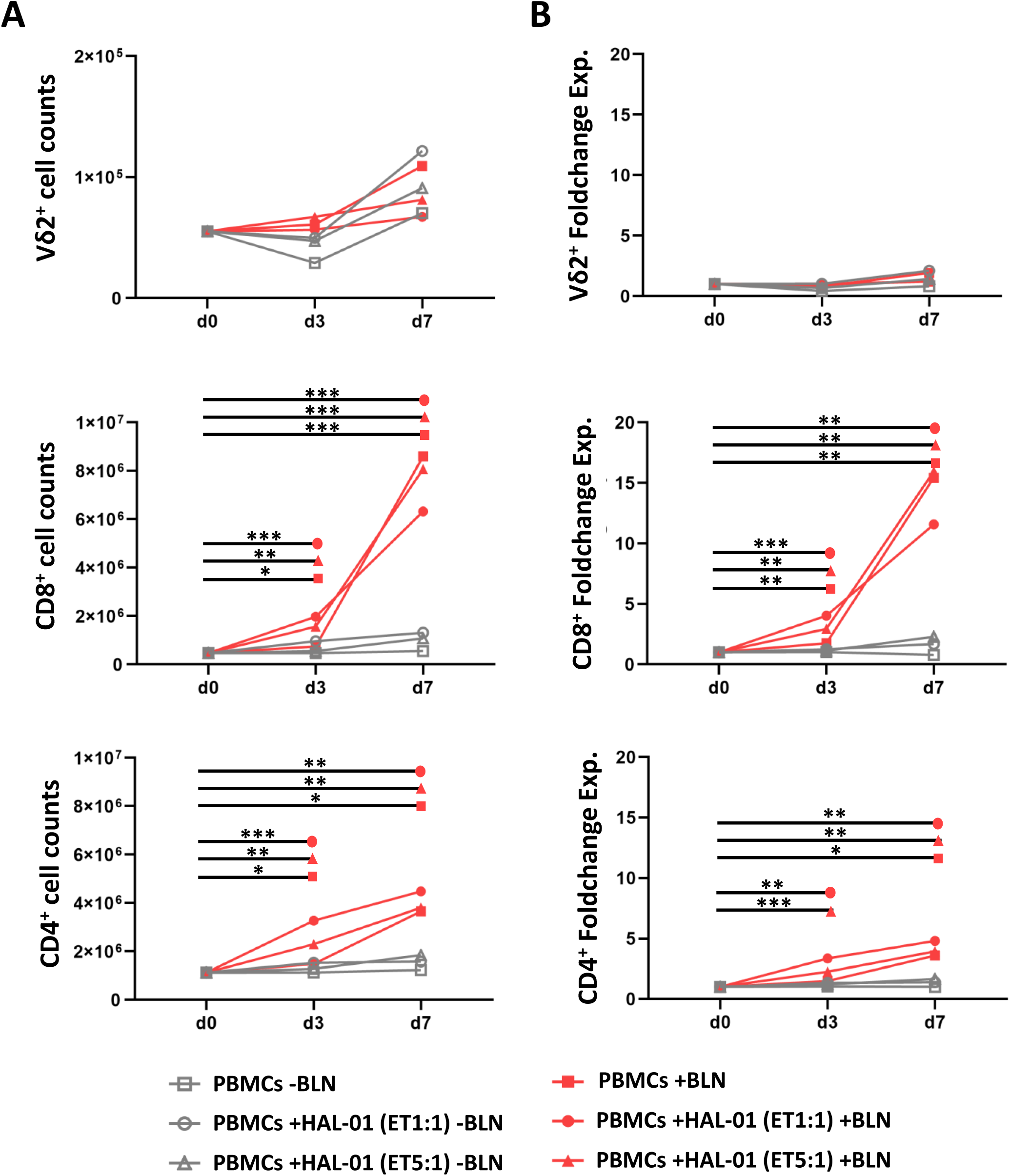
BLN-induced expansion of CD8 αβ but not γδ T cells in ex vivo cultures. A: Absolute cell counts of Vδ2^+^ CD8^+^ and CD4^+^ T cells after three and seven days of PBMC stimulation with BLN (red symbols, continuous lines) or without BLN (grey symbols, dashed lines). Cultures without additional malignant B cells are marked as squares (containing autologous B cells), cultures with low tumor load (E:T 5:1) as triangles and medium tumor load (E:T 1:1) as circles. B: Foldchange expansion of Vδ2^+^, CD8^+^ and CD4^+^ T cells after three and seven days.

### scRNA–seq reveals strong activation/exhaustion programs in αβ T cells

To compare transcriptional profiles of αβ and γδ T cells following BLN stimulation at single-cell resolution, we performed single-cell RNA sequencing using the BD Rhapsody™ platform with a targeted human T-cell panel focusing on key genes involved in T-cell responses. PBMCs from a healthy donor (containing autologous B cells) were cultured for 3 days with or without BLN (20 ng/mL). Subsequently, CD3⁺ T cells were negatively isolated for sequencing. In parallel, immunophenotyping was performed using AbSeq reagents and conventional flow cytometry to support cluster identification and validate phenotypic states (Figure 6B). After quality control and removing non-viable cells, 6872 high-quality T cells were included in the analysis. Dimensionality reduction and Seurat-based clustering on UMAP space identified 14 initial clusters, which were manually curated and consolidated into eight main functional clusters (Figure 6C, Supplemental Figure 5). UMAP projection showed a clear separation between BLN-treated (red, *3,576 cells*) and untreated (grey *3,296 cells*) conditions, indicating major shifts in transcriptomic profiles upon BLN stimulation. While γδ T cells (mainly clustering in groups 6 and 13) showed minimal transcriptional changes following BLN exposure, αβ T cells, particularly CD8⁺ T cells, underwent pronounced transcriptional changes. BLN stimulation led to increased expression of genes associated with proliferation and effector function, as well as exhaustion markers including TIGIT, TIM-3 (HAVCR2), LAG3, and CD160 (Figure 6C–E; Supplemental Figure 5). In contrast, γδ T-cell frequencies remained largely stable, consistent with the phenotypic data and aligned with in vitro, with lack of γδ T cell expansion from PBMCs at day 3 as shown in Figure 5. Of note again, also on transcript level, exhaustion marker TIGIT was not expressed on γδ T cells following BLN exposure^54^.

**Figure 6.**
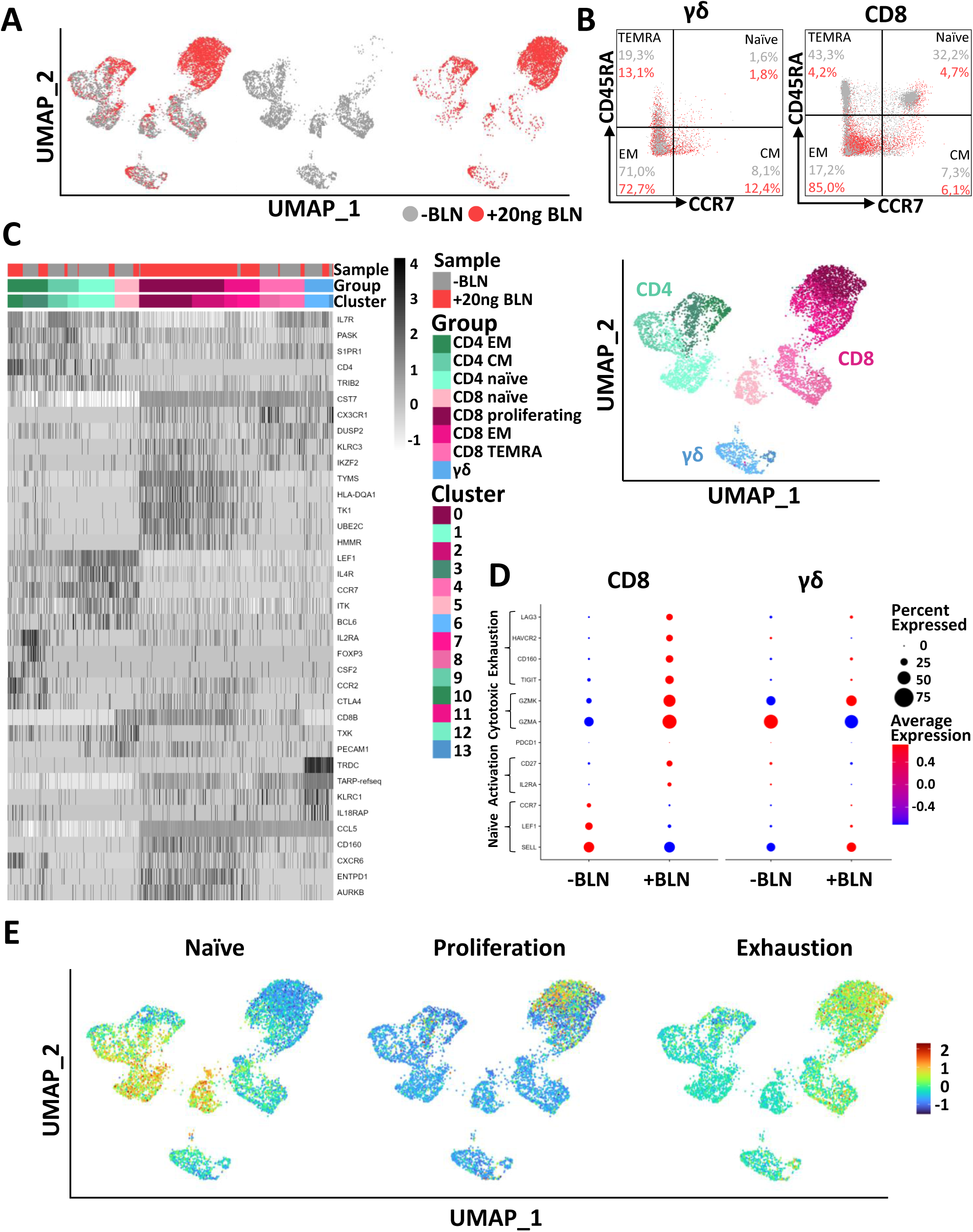
scRNA-seq reveals strong activation/exhaustion programs in αβ T cells. (A) UMAP plots of Seurat clusters from all CD3⁺ cells: combined map (left; without BLN = grey, with 20 ng/mL BLN = red), and condition-specific maps (middle = without BLN; right = with BLN). (B) T-cell subsets (naïve, CM, EM, TEMRA) distinguished by CD45RA and CCR7 expression on γδ and CD8⁺ T cells by flow cytometry. (C) Heatmap and UMAP of 14 original and 8 manually curated clusters representing CD4⁺, CD8⁺, and γδ T cells. (D) Expression changes of selected genes in CD8⁺ and γδ T cells (dot size = % of cells expressing; color = average expression). (E) UMAP highlighting genes representing naïve, proliferative, and exhausted states (gene sets listed in Supplemental Table 3).

## Discussion

BLN is a highly potent bispecific T-cell engager directing T-cell-mediated cytotoxicity toward CD19-expressing malignant blasts and normal B cells. In this study, we dissected BLN-mediated cytotoxic mechanisms by directly comparing resting and activated CD3-expressing conventional αβ T cells (CD4⁺ and CD8⁺) and non-conventional γδ T cells, to evaluate their cytotoxic functions and phenotypic shifts following BLN stimulation.

Here we show that BLN efficiently redirected cytotoxicity toward CD19⁺ targets across αβ and γδ T-cell subsets. However, freshly isolated γδ T cells demonstrated reduced BLN-mediated cytotoxicity against CD19⁺ tumor cells compared to αβ T cells. Because BLN binds CD3ε rather than the TCR variable domains, differences between αβ and γδ T cells under BLN likely reflect CD3 epitope/clone biology and CD3 glycosylation. In addition, lineage-specific signaling might also play a role, as antigen stimulation of the γδ TCR itself does not trigger CD3 conformational change (CD3 CC), in contrast to αβ T cells.^29,55–57^ In human Vγ9Vδ2 γδ T cells, effects of anti-CD3 clones differ markedly: UCHT1 clone enforces CD3 CC and strongly augments γδ cytotoxicity, whereas OKT3 clone is comparatively weak unless valency is increased or CD3 is deglycosylated.^55^ In BLN CD3 scFv derives from murine L2K 07 clone. Some older reviews have mentioned TR66 as the parental clone of anti–CD3 for BLN,^58^ but more recent primary sources identify L2K–07 as the basis.^59–61^ However, direct functional comparison of clone L2K 07 with UCHT1 and OKT3 in γδ T cells are still missing.

Among αβ T cells, CD8⁺ T cells mount a stronger response to BLN than CD4⁺ T cells, consistent with prior reports showing superior cytotoxic potency of CD8⁺ T cells in the presence of BLN^53^. Collectively, these findings support notion that, in vivo, CD8⁺ T cells are the principal contributors to therapeutic responses to BLN. Given the low γδ T-cell frequencies in blood, these cells are readily outcompeted by αβ cells in mixed cultures, which likely contributed to the minimal γδ expansion observed ex vivo.

In contrast, γδ T-cell lines, activated and expanded ex vivo using Zole, which selectively expands the Vγ9Vδ2 γδ T-cell subset, exhibited comparable cytotoxic efficacy to PHA-expanded αβ T cells. Across experiments, antigen density and tumor burden emerged as dominant modulators of potency, and BLN dosing further tuned γδ T-cell activity. Both Zole–expanded γδ and PHA–expanded αβ T cells cleared CD19⁺ targets under low tumor burden, whereas at high tumor burden αβ T cells quickly eliminated CD19^+^ target and whereas γδ T-cell-killing benefited from higher BLN concentration over time^29^. Other T–cell engager designs most likely constructed with the same CD3–binder lineage as BLN^62^ (e.g., HER2×CD3) have shown potent γδ–mediated cytotoxicity in vitro. These experiments used much higher bispecific doses and higher effector–to–target ratios compared to condition here^57,62^. This highlights that our results do not indicate a limitation of γδ cells under CD3-engagers; they show potency of BLN-stimulated γδ T cells and that αβ cells respond more robustly to BLN than γδ cells—at the cost of a higher risk of overt activation. We showed that BLN triggered sustained surface CD3 down–modulation in αβ T cells but not in γδ T cells, particularly at high target-load and was accompanied by consistently higher sFas-L release in αβ cultures. High disease burden amplifies BLN-driven synapse frequency and CD3 signaling^40^, predisposing αβ T cells to early AICD. This level of TCR–CD3 engagement is known to induce CD3 down modulation (internalization and ζ chain degradation) and to trigger Fas/FasL mediated AICD on re-stimulation^45^. Our observation of higher sFas-L secretion together with early CD3 down–modulation in αβ therefore supports early AICD in CD8^+^ αβ T cells. Further, the preferential CD8 T-cell expansion at lower target load from co-cultures of activated T cells and mixed PBMC cultures supports this hypothesis. Clinically, the same principle underlies superior BLN outcomes at lower leukemia burden, the use of debulking, and BLN’s positioning in MRD settings and often deploy it as a bridge to allogeneic transplant^8,12,63,64^. This altogether suggests that high CD19 load accelerates αβ activation but precipitates early AICD and limits CD8⁺ expansion, whereas ZOL–expanded Vγ9Vδ2 γδ T cells retain CD3 and probably resist AICD.

Importantly day 14 Zole-expanded γδ T cells displayed an effector phenotype with low expression of inhibitory checkpoints. Conversely, PHA–expanded αβ T cells underwent rapid activation and differentiation from naïve to EM/TEMRA. Single–cell RNA–seq mirrored these findings: αβ cells acquired proliferative and activation/exhaustion programs, whereas γδ cells showed comparatively modest transcriptional shifts.

In our assays, γδ cultures while exerting cytotoxic effector functions, produced lower IFNγ and cytotoxic–granule release than αβ cultures and did not expand under BLN stimulation. Because cytokine release syndrome (CRS) and Immune Effector Cell-Associated Neurotoxicity Syndrome (ICANS) are linked to systemic cytokine surges and widespread T–cell activation during CD3–engager therapy, these features predict lower inflammatory risk with γδ–based strategies. Therefore, γδ adoptive transfer alongside BLN therapy may deliver additional tumor control with a reduced probability of high–grade CRS/ICANS.

Zole expansion protocols have become a standard approach for generating clinical-grade γδ T-cell products^39^. Several ongoing clinical trials^19^ are evaluating Zole-expanded γδ T cells, both as monotherapy and in combination with BLN, to enhance treatment efficacy and improve T-cell fitness, especially in the context of heavily pretreated patients undergoing chemotherapy. A key advantage of γδ T cells in adoptive transfer settings is their innate-like, non-alloreactive profile, eliminating concerns related to GvHD, thus making them particularly attractive for allogeneic and off-the-shelf therapy. This therapy should be prioritizing γδ + BLN in low–burden/MRD disease and in patients with suboptimal αβ fitness, where γδ T-cells can potentiate cytotoxicity without amplifying inflammatory risk, as supported by preclinical CD19-BiTE and Vγ9Vδ2 γδ T-cell synergy^34^. Additionally, optimizing γδ T-cell expansion protocols—for instance, incorporating cytokines IL-2, IL-15, and vitamin C, has been shown to significantly boost cytotoxic function and proliferation while reducing apoptosis, suggesting a promising direction for future clinical applications. This approach with 414 expanded γδ T-cell infusions into 132 cancer patients has been shown to be clinically safe without significant adverse effects and prolonged the survival of late-stage cancer patients who received ≥5 cell infusions in a small cohort of 8 liver and 10 lung cancer patients^32^.

In conclusion, our findings emphasize the therapeutic potential of γδ T cells in adoptive T-cell therapies, particularly when expanded ex vivo with Zole.^19,39,44^ Their stable cytotoxic functionality, intrinsic resistance to AICD, and lack of alloreactivity provide significant advantages in the design of robust, clinically effective, adoptive immunotherapy protocols.

Future clinical studies are warranted to evaluate the clinical efficacy and long-term benefits of Zole-expanded γδ T-cell products alone or in combination with BLN and other immune modulators, to harness their full therapeutic potential and improve outcomes for patients with CD19-expressing malignancies.

## Acknowledgments

We dedicate this work to the memory of Dr. Marcus Lettau, whose scientific insight, commitment, and collegial spirit were central to the conception of this study. Together with the corresponding author, he designed the project and initiated key experiments that formed the foundation of the present work. His enthusiasm for translational immunology and mentorship continues to inspire our research.

## Authorship Contributions

MK, NN, SB performed experiments and analyzed data. MK performed statistical analysis and prepared first draft of the results and materials and methods. GC and ML designed and conceptualized the project. HT, CP, DK, OJ, DW, and HHO supported in vitro culture experiments and provided expert assistance. AL helped with multiparametric flow cytometry analysis. ML and TV helped with quantification of CD19 on tumor cell lines. GC, CBD, MB, DK, DW, HHO and OJ helped with data interpretation. GC finalized manuscript. All authors reviewed the manuscript and approved its final version.

## References

1. Liu D, Zhao J, Song Y, Luo X, Yang T. Clinical trial update on bispecific antibodies, antibody-drug conjugates, and antibody-containing regimens for acute lymphoblastic leukemia. J Hematol Oncol 2019;12:15, DOI: 10.1186/s13045-019-0703-z.

2. Chitadze G, Laqua A, Lettau M, Baldus CD, Bruggemann M. Bispecific antibodies in acute lymphoblastic leukemia therapy. Expert Rev Hematol 2020;13:1211–33, DOI: 10.1080/17474086.2020.1831380.

3. Dinmohamed AG, Szabo A, van der Mark M, et al. Improved survival in adult patients with acute lymphoblastic leukemia in the Netherlands: a population-based study on treatment, trial participation and survival. Leukemia 2016;30:310–7, DOI: 10.1038/leu.2015.230.

4. Fredman D, Moshe Y, Wolach O, et al. Evaluating outcomes of adult patients with acute lymphoblastic leukemia and lymphoblastic lymphoma treated on the GMALL 07/2003 protocol. Ann Hematol 2022;101:581–93, DOI: 10.1007/s00277-021-04738-y.

5. Möricke A, Zimmermann M, Valsecchi MG, et al. Dexamethasone vs prednisone in induction treatment of pediatric ALL: results of the randomized trial AIEOP-BFM ALL 2000. Blood 2016;127:2101–12, DOI: 10.1182/blood-2015-09-670729.

6. Baden D, Wolgast N, Altrock PM, et al. Epidemiology, survival, and treatment of acute myeloid and lymphoblastic leukaemia in Germany: a nationwide population-based registry analysis. The Lancet Regional Health - Europe 2025;59:101503, DOI: 10.1016/j.lanepe.2025.101503.

7. Bargou R, Leo E, Zugmaier G, et al. Tumor regression in cancer patients by very low doses of a T cell-engaging antibody. Science 2008;321:974–7, DOI: 10.1126/science.1158545.

8. Kantarjian H, Stein A, Gokbuget N, et al. Blinatumomab versus Chemotherapy for Advanced Acute Lymphoblastic Leukemia. N Engl J Med 2017;376:836–47, DOI: 10.1056/NEJMoa1609783.

9. von Stackelberg A, Locatelli F, Zugmaier G, et al. Phase I/Phase II Study of Blinatumomab in Pediatric Patients With Relapsed/Refractory Acute Lymphoblastic Leukemia. J Clin Oncol 2016;34:4381–9, DOI: 10.1200/JCO.2016.67.3301.

10. Gupta S, Rau RE, Kairalla JA, et al. Blinatumomab in Standard-Risk B-Cell Acute Lymphoblastic Leukemia in Children. N Engl J Med 2025;392:875–91, DOI: 10.1056/NEJMoa2411680.

11. Litzow MR, Sun Z, Mattison RJ, et al. Blinatumomab for MRD-Negative Acute Lymphoblastic Leukemia in Adults. N Engl J Med 2024;391:320–33, DOI: 10.1056/NEJMoa2312948.

12. Gokbuget N, Dombret H, Bonifacio M, et al. Blinatumomab for minimal residual disease in adults with B-cell precursor acute lymphoblastic leukemia. Blood 2018;131:1522–31, DOI: 10.1182/blood-2017-08-798322.

13. Topp MS, Gokbuget N, Stein AS, et al. Safety and activity of blinatumomab for adult patients with relapsed or refractory B-precursor acute lymphoblastic leukaemia: a multicentre, single-arm, phase 2 study. Lancet Oncol 2015;16:57–66, DOI: 10.1016/S1470-2045(14)71170-2.

14. Locatelli F, Shah B, Thomas T, et al. Incidence of CD19-negative relapse after CD19-targeted immunotherapy in R/R BCP acute lymphoblastic leukemia: a review. Leuk Lymphoma 2023;64:1615–33, DOI: 10.1080/10428194.2023.2232496.

15. Das RK, O’Connor RS, Grupp SA, Barrett DM. Lingering effects of chemotherapy on mature T cells impair proliferation. Blood Adv 2020;4:4653–64, DOI: 10.1182/bloodadvances.2020001797.

16. Philipp N, Kazerani M, Nicholls A, et al. T-cell exhaustion induced by continuous bispecific molecule exposure is ameliorated by treatment-free intervals. Blood 2022;140:1104–18, DOI: 10.1182/blood.2022015956.

17. Hijazi Y, Klinger M, Kratzer A, et al. Pharmacokinetic and Pharmacodynamic Relationship of Blinatumomab in Patients with Non-Hodgkin Lymphoma. Curr Clin Pharmacol 2018;13:55–64, DOI: 10.2174/1574884713666180518102514.

18. Kabelitz D, Serrano R, Kouakanou L, Peters C, Kalyan S. Cancer immunotherapy with gammadelta T cells: many paths ahead of us. Cell Mol Immunol 2020;17:925–39, DOI: 10.1038/s41423-020-0504-x.

19. Hayday A, Dechanet-Merville J, Rossjohn J, Silva-Santos B. Cancer immunotherapy by gammadelta T cells. Science 2024;386:eabq7248, DOI: 10.1126/science.abq7248.

20. Gober HJ, Kistowska M, Angman L, Jeno P, Mori L, De Libero G. Human T cell receptor gammadelta cells recognize endogenous mevalonate metabolites in tumor cells. J Exp Med 2003;197:163–8, DOI: 10.1084/jem.20021500.

21. Willcox CR, Mohammed F, Willcox BE. The distinct MHC-unrestricted immunobiology of innate-like and adaptive-like human gammadelta T cell subsets-Nature’s CAR-T cells. Immunol Rev 2020;298:25–46, DOI: 10.1111/imr.12928.

22. Gaballa A, Arruda LCM, Uhlin M. Gamma delta T-cell reconstitution after allogeneic HCT: A platform for cell therapy. Front Immunol 2022;13:971709, DOI: 10.3389/fimmu.2022.971709.

23. Godder KT, Henslee-Downey PJ, Mehta J, et al. Long term disease-free survival in acute leukemia patients recovering with increased gammadelta T cells after partially mismatched related donor bone marrow transplantation. Bone Marrow Transplant 2007;39:751–7, DOI: 10.1038/sj.bmt.1705650.

24. Esin S, Shigematsu M, Nagai S, Eklund A, Wigzell H, Grunewald J. Different percentages of peripheral blood gamma delta + T cells in healthy individuals from different areas of the world. Scand J Immunol 1996;43:593–6, DOI: 10.1046/j.1365-3083.1996.d01-79.x.

25. Davey MS, Willcox CR, Hunter S, et al. The human Vdelta2(+) T-cell compartment comprises distinct innate-like Vgamma9(+) and adaptive Vgamma9(-) subsets. Nat Commun 2018;9:1760, DOI: 10.1038/s41467-018-04076-0.

26. Roelofs AJ, Jauhiainen M, Monkkonen H, Rogers MJ, Monkkonen J, Thompson K. Peripheral blood monocytes are responsible for gammadelta T cell activation induced by zoledronic acid through accumulation of IPP/DMAPP. Br J Haematol 2009;144:245–50, DOI: 10.1111/j.1365-2141.2008.07435.x.

27. Kondo M, Sakuta K, Noguchi A, et al. Zoledronate facilitates large-scale ex vivo expansion of functional gammadelta T cells from cancer patients for use in adoptive immunotherapy. Cytotherapy 2008;10:842–56, DOI: 10.1080/14653240802419328.

28. Harly C, Guillaume Y, Nedellec S, et al. Key implication of CD277/butyrophilin-3 (BTN3A) in cellular stress sensing by a major human gammadelta T-cell subset. Blood 2012;120:2269–79, DOI: 10.1182/blood-2012-05-430470.

29. Rigau M, Ostrouska S, Fulford TS, et al. Butyrophilin 2A1 is essential for phosphoantigen reactivity by γδ T cells. Science 2020;367:eaay5516, DOI: doi:10.1126/science.aay5516.

30. Brandes M, Willimann K, Bioley G, et al. Cross-presenting human gammadelta T cells induce robust CD8+ alphabeta T cell responses. Proc Natl Acad Sci U S A 2009;106:2307–12, DOI: 10.1073/pnas.0810059106.

31. Holmen Olofsson G, Idorn M, Carnaz Simoes AM, et al. Vgamma9Vdelta2 T Cells Concurrently Kill Cancer Cells and Cross-Present Tumor Antigens. Front Immunol 2021;12:645131, DOI: 10.3389/fimmu.2021.645131.

32. Xu Y, Xiang Z, Alnaggar M, et al. Allogeneic Vgamma9Vdelta2 T-cell immunotherapy exhibits promising clinical safety and prolongs the survival of patients with late-stage lung or liver cancer. Cell Mol Immunol 2021;18:427–39, DOI: 10.1038/s41423-020-0515-7.

33. Kelm M, Darzentas F, Darzentas N, et al. Dominant T-cell Receptor Delta Rearrangements in B-cell Precursor Acute Lymphoblastic Leukemia: Leukemic Markers or Physiological gammadelta T Repertoire? Hemasphere 2023;7:e948, DOI: 10.1097/HS9.0000000000000948.

34. Chen YH, Wang Y, Liao CH, Hsu SC. The potential of adoptive transfer of gamma9delta2 T cells to enhance blinatumomab’s antitumor activity against B-cell malignancy. Sci Rep 2021;11:12398, DOI: 10.1038/s41598-021-91784-1.

35. You H, Zhu H, Zhao Y, Guo J, Gao Q. TIGIT-expressing zoledronate-specific gammadelta T cells display enhanced antitumor activity. J Leukoc Biol 2022;112:1691–700, DOI: 10.1002/JLB.5MA0822-759R.

36. Chen D, Guo Y, Jiang J, et al. gammadelta T cell exhaustion: Opportunities for intervention. J Leukoc Biol 2022;112:1669–76, DOI: 10.1002/JLB.5MR0722-777R.

37. Bender A, Kabelitz D. CD4-CD8-human T cells: phenotypic heterogeneity and activation requirements of freshly isolated “double-negative” T cells. Cell Immunol 1990;128:542–54, DOI: 10.1016/0008-8749(90)90047-u.

38. Katzen D, Chu E, Terhost C, et al. Mechanisms of human T cell response to mitogens: IL 2 induces IL 2 receptor expression and proliferation but not IL 2 synthesis in PHA-stimulated T cells. J Immunol 1985;135:1840–5.

39. Peters C, Kouakanou L, Oberg HH, Wesch D, Kabelitz D. In vitro expansion of Vgamma9Vdelta2 T cells for immunotherapy. Methods Enzymol 2020;631:223–37, DOI: 10.1016/bs.mie.2019.07.019.

40. Liu C, Zhou J, Kudlacek S, Qi T, Dunlap T, Cao Y. Population dynamics of immunological synapse formation induced by bispecific T cell engagers predict clinical pharmacodynamics and treatment resistance. Elife 2023;12, DOI: 10.7554/eLife.83659.

41. Hao Y, Stuart T, Kowalski MH, et al. Dictionary learning for integrative, multimodal and scalable single-cell analysis. Nat Biotechnol 2024;42:293–304, DOI: 10.1038/s41587-023-01767-y.

42. Saini RV, Wilson C, Finn MW, Wang T, Krensky AM, Clayberger C. Granulysin delivered by cytotoxic cells damages endoplasmic reticulum and activates caspase-7 in target cells. J Immunol 2011;186:3497–504, DOI: 10.4049/jimmunol.1003409.

43. Seder RA, Ahmed R. Similarities and differences in CD4+ and CD8+ effector and memory T cell generation. Nat Immunol 2003;4:835–42, DOI: 10.1038/ni969.

44. Kondo M, Izumi T, Fujieda N, et al. Expansion of human peripheral blood gammadelta T cells using zoledronate. J Vis Exp 2011, DOI: 10.3791/3182.

45. Green DR, Droin N, Pinkoski M. Activation-induced cell death in T cells. Immunol Rev 2003;193:70–81, DOI: 10.1034/j.1600-065x.2003.00051.x.

46. Ebsen H, Lettau M, Kabelitz D, Janssen O. Subcellular localization and activation of ADAM proteases in the context of FasL shedding in T lymphocytes. Mol Immunol 2015;65:416–28, DOI: 10.1016/j.molimm.2015.02.008.

47. Dreier T, Lorenczewski G, Brandl C, et al. Extremely potent, rapid and costimulation-independent cytotoxic T-cell response against lymphoma cells catalyzed by a single-chain bispecific antibody. Int J Cancer 2002;100:690–7, DOI: 10.1002/ijc.10557.

48. Baeuerle PA, Kufer P, Bargou R. BiTE: Teaching antibodies to engage T-cells for cancer therapy. Curr Opin Mol Ther 2009;11:22–30.

49. Hoffmann P, Hofmeister R, Brischwein K, et al. Serial killing of tumor cells by cytotoxic T cells redirected with a CD19-/CD3-bispecific single-chain antibody construct. Int J Cancer 2005;115:98–104, DOI: 10.1002/ijc.20908.

50. Zhu M, Kratzer A, Johnson J, et al. Blinatumomab Pharmacodynamics and Exposure-Response Relationships in Relapsed/Refractory Acute Lymphoblastic Leukemia. J Clin Pharmacol 2018;58:168–79, DOI: 10.1002/jcph.1006.

51. Valitutti S, Muller S, Salio M, Lanzavecchia A. Degradation of T cell receptor (TCR)-CD3-zeta complexes after antigenic stimulation. J Exp Med 1997;185:1859–64, DOI: 10.1084/jem.185.10.1859.

52. Suda T, Hashimoto H, Tanaka M, Ochi T, Nagata S. Membrane Fas ligand kills human peripheral blood T lymphocytes, and soluble Fas ligand blocks the killing. J Exp Med 1997;186:2045–50, DOI: 10.1084/jem.186.12.2045.

53. Klinger M, Brandl C, Zugmaier G, et al. Immunopharmacologic response of patients with B-lineage acute lymphoblastic leukemia to continuous infusion of T cell-engaging CD19/CD3-bispecific BiTE antibody blinatumomab. Blood 2012;119:6226–33, DOI: 10.1182/blood-2012-01-400515.

54. Rancan C, Arias-Badia M, Dogra P, et al. Exhausted intratumoral Vdelta2(-) gammadelta T cells in human kidney cancer retain effector function. Nat Immunol 2023;24:612–24, DOI: 10.1038/s41590-023-01448-7.

55. Dopfer EP, Hartl FA, Oberg HH, et al. The CD3 conformational change in the gammadelta T cell receptor is not triggered by antigens but can be enforced to enhance tumor killing. Cell Rep 2014;7:1704–15, DOI: 10.1016/j.celrep.2014.04.049.

56. Oberg HH, Kellner C, Gonnermann D, et al. gammadelta T cell activation by bispecific antibodies. Cell Immunol 2015;296:41–9, DOI: 10.1016/j.cellimm.2015.04.009.

57. Alarcon B, De Vries J, Pettey C, et al. The T-cell receptor gamma chain-CD3 complex: implication in the cytotoxic activity of a CD3+ CD4-CD8-human natural killer clone. Proc Natl Acad Sci U S A 1987;84:3861–5, DOI: 10.1073/pnas.84.11.3861.

58. Burt R, Warcel D, Fielding AK. Blinatumomab, a bispecific B-cell and T-cell engaging antibody, in the treatment of B-cell malignancies. Hum Vaccin Immunother 2019;15:594–602, DOI: 10.1080/21645515.2018.1540828.

59. Jacobs N, Mazzoni A, Mezzanzanica D, et al. Efficiency of T cell triggering by anti-CD3 monoclonal antibodies (mAb) with potential usefulness in bispecific mAb generation. Cancer Immunol Immunother 1997;44:257–64, DOI: 10.1007/s002620050381.

60. Mocquot P, Mossazadeh Y, Lapierre L, Pineau F, Despas F. The pharmacology of blinatumomab: state of the art on pharmacodynamics, pharmacokinetics, adverse drug reactions and evaluation in clinical trials. J Clin Pharm Ther 2022;47:1337–51, DOI: 10.1111/jcpt.13741.

61. Blinatumomab (AMG 103, BLINCYTO®). https://dctd.cancer.gov, 2025.

62. Oberg HH, Peipp M, Kellner C, et al. Novel bispecific antibodies increase gammadelta T-cell cytotoxicity against pancreatic cancer cells. Cancer Res 2014;74:1349–60, DOI: 10.1158/0008-5472.CAN-13-0675.

63. Queudeville M, Stein AS, Locatelli F, et al. Low leukemia burden improves blinatumomab efficacy in patients with relapsed/refractory B-cell acute lymphoblastic leukemia. Cancer 2023;129:1384–93, DOI: 10.1002/cncr.34667.

64. Bonifacio M, Papayannidis C, Lussana F, et al. Real-World Multicenter Experience in Tumor Debulking Prior to Blinatumomab Administration in Adult Patients With Relapsed/Refractory B-Cell Precursor Acute Lymphoblastic Leukemia. Front Oncol 2021;11:804714, DOI: 10.3389/fonc.2021.804714.

