## Supplemental_Figures.pdf for "Blinatumomab-driven T-cell activation in αβ and γδ T-cell subsets: Insights from in vitro assays§"

PHA Gating Strategy d0

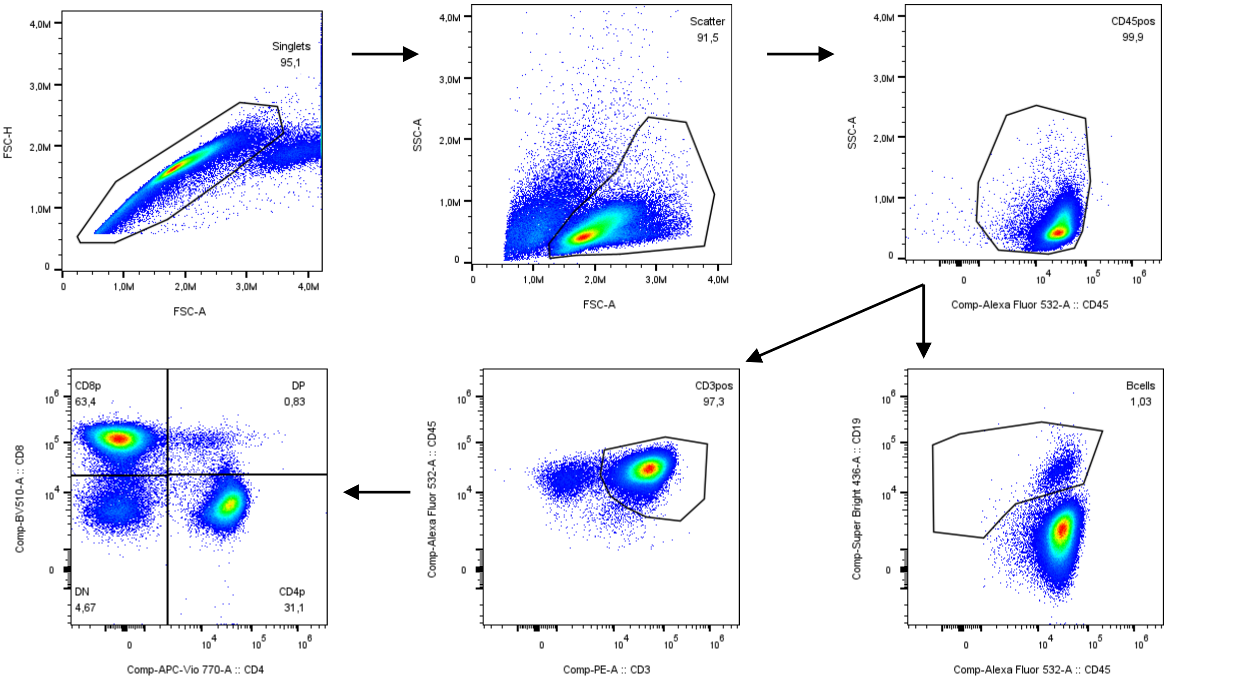


Zole Gating Strategy d0

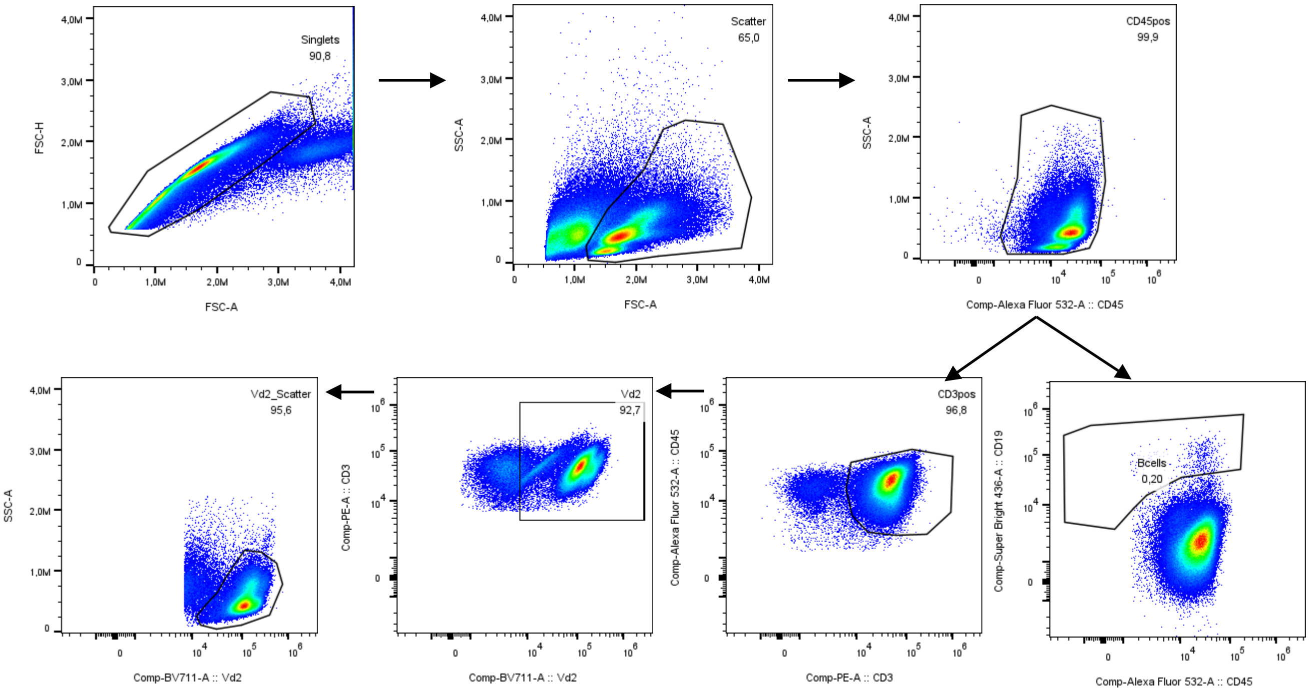


Supplemental Figure 1: Gating Strategy of PHA-expanded αβ and Zole expanded γδ T cells.

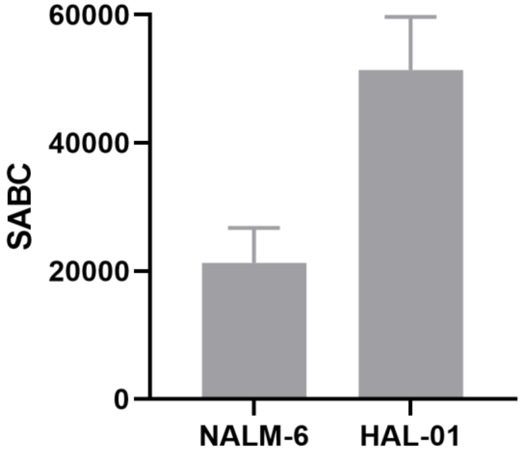


Supplemental Figure 2: Quantitative determination of CD19-antigen expression on B-cell surface of HAL-01 and NALM-6 cell lines as analyzed by flow cytometry using the purified anti-human CD19 antibody (clone HIB19) and QIFIKIT®. Shown are the median specific antibody binding capacity (SABC) values + 95% CI of four experiments.

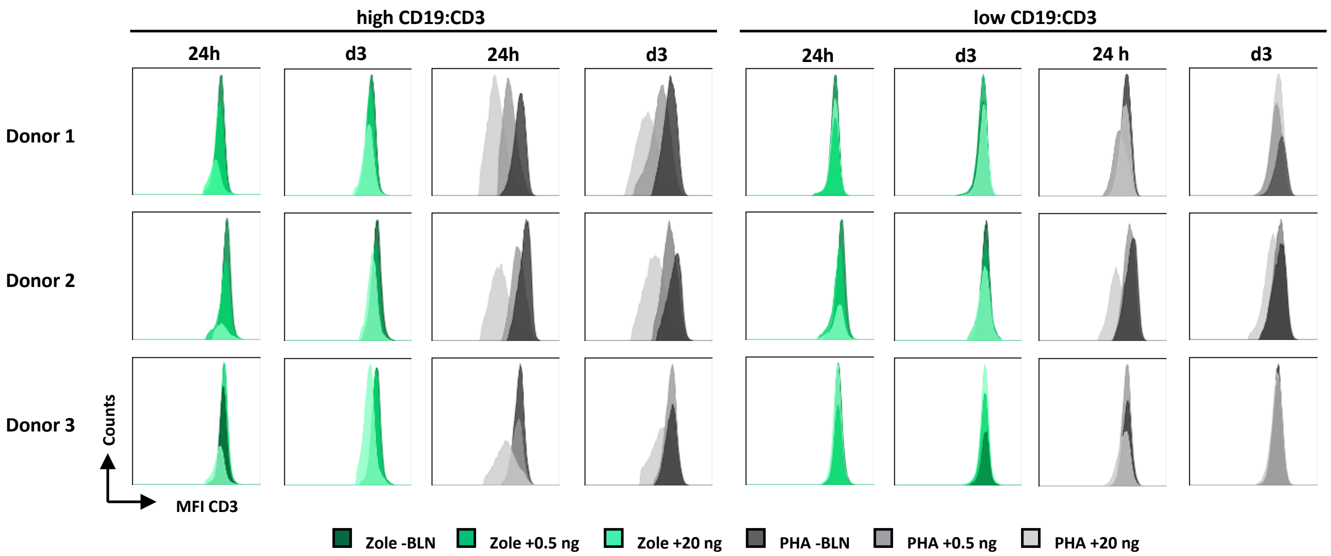


Supplemental Figure 3. CD3 median fluorescence intensity (MFI) under different BLN concentrations (-BLN, 0.5ng/ml, 20ng/ml) and varying CD19-expressing tumor load per single T cell (high: CD19:CD3 load (5:1) and low CD19:CD3 load (1:5) after 24h and three days (d3) of all three donors.

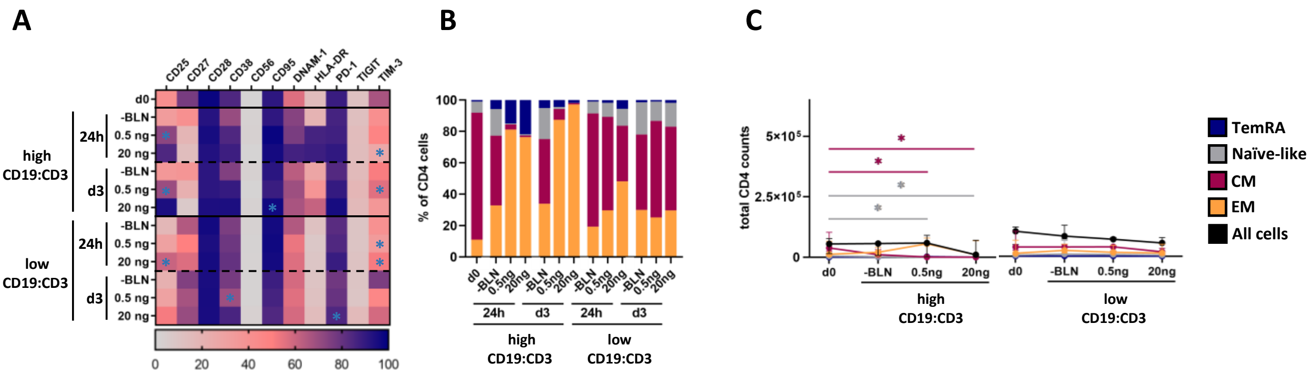


Supplemental Figure 4. Phenotypic changes in CD4⁺ αβ T cells under varying concetrations of BLN and target effector ratios. between Zole- and PHA-cultures.

A: Median Marker Expression on PHA-expanded CD4⁺ T cells of three healthy donors. B: Median values of EM, CM, Naïve and TEMRA cells of CD4⁺ T cells. B: Absolute cell counts of EM, CM, Naïve and TEMRA cells of CD4⁺ αβ T cells at day 0 and day 3. Error bars representing Median value with 95% confidence interval. Stars representing significant changes between baseline and BLN therapy at day 1 and day 3.

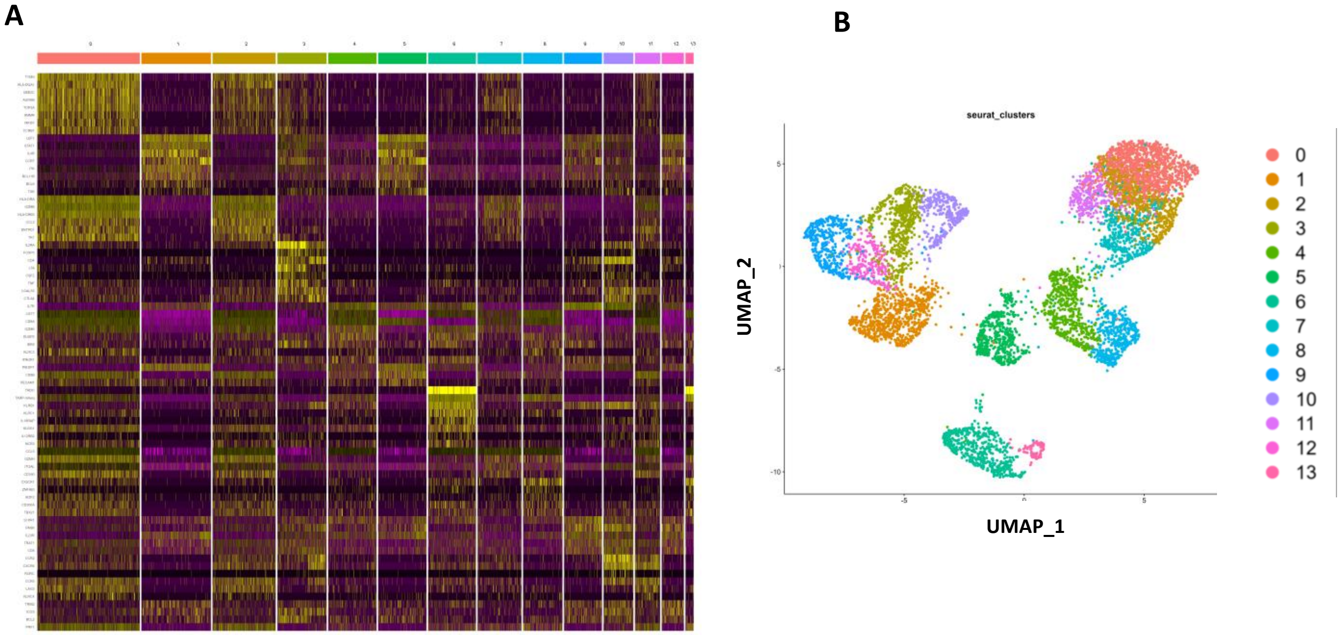


Supplemental Figure 5: Single-cell RNA-sequencing of in vitro stimulated PBMC. A. Seurat clustering identified 14 distinct population that are mapped om UMAP 2D (B) space as follows: 0: CD8+. proliferating; 1: CD4+ naïve; 2: CD8+ proliferating; 3: CD4+ EM; 4: CD8+ TEMRA; 5: CD8+ naive; 6: γδ+; 7: CD8+ EM; 8: CD8+ TEMRA; 9: CM; 10: CD4+ EM; 11: CD8+ EM; 12: CD4+ CM; 13: γδ+
